# Repeated stimulation of the HPA axis alters white blood cell counts without increasing oxidative stress or inflammatory cytokines in fasting elephant seal pups

**DOI:** 10.1101/2021.03.31.437957

**Authors:** David C. Ensminger, Daniel E. Crocker, Emily K. Lam, N. Allen Kaitlin, José Pablo Vázquez-Medina

## Abstract

The hypothalamic-pituitary-adrenal (HPA) axis controls the release of glucocorticoids, which regulate immune and inflammatory function by modulating cytokines, white blood cells (WBCs), and oxidative stress via glucocorticoid receptor (GR) signaling. Although the response to HPA activation is well characterized in many species, little is known about the impacts of HPA activation during extreme physiological conditions in marine mammals. Hence, we challenged 18 simultaneously fasting and developing elephant seal pups with daily intramuscular injections of adrenocorticotropin (ACTH), a GR antagonist (RU486), or a combination (ACTH+RU486) for four days (4d). We collected blood at baseline, two hours (2h), and 4d after the beginning of treatment. ACTH and ACTH+RU486 elevated serum aldosterone and cortisol at 2h, with effects diminishing at 4d. RU486 alone induced a compensatory increase in aldosterone, but not cortisol, at 4d. ACTH decreased neutrophils at 2h while decreasing lymphocytes and increasing neutrophil:lymphocyte ratio at 4d. These effects were abolished by RU486. Despite alterations in WBCs, there was no effect of ACTH or RU486 on transforming growth factor-β or interleukin-6 levels; however, both cytokines decreased with the 4-d fasting progression. Similarly, ACTH did not impact protein oxidation, lipid peroxidation, or antioxidant enzymes, but plasma isoprostanes and catalase activity decreased while glutathione peroxidase increased with fasting progression. These data demonstrate differential acute (2h) and chronic (4d) modulatory effects of HPA activation on WBCs and that the chronic effect is mediated, at least in part, by GR. These results also underscore elephant seals’ resistance to potential oxidative stress derived from repeated HPA activation.

**Summary statement:** Many species experience oxidative stress and inflammation after repeated activation of the hypothalamic-pituitary-adrenal axis. We show that simultaneously fasting and developing elephant seals are resistant to repeated hypothalamic-pituitary-adrenal axis activation.

## 1. Introduction

The hypothalamic-pituitary-adrenal (HPA) axis facilitates organismal responses to metabolic perturbations and environmental stressors (Sapolsky et al., 2000; Tsigos and Chrousos, 2002). Upon activation of the HPA axis, the hypothalamus secretes corticotropin-releasing hormone, which stimulates the anterior pituitary to release adrenocorticotropin (ACTH; Plotsky et al., 1989). ACTH then acts on the adrenal cortex to induce the secretion of corticosteroids including the glucocorticoid cortisol and the mineralocorticoid aldosterone (Haning and Tait, 1970). Cortisol and aldosterone promote energy mobilization and osmotic balance via signaling mechanisms involving glucocorticoid (GR) and mineralocorticoid receptors (MacDougall-Shackleton et al., 2019). Thus, adrenocorticosteroids impact many physiological processes including lipolysis, immune function, and oxidative stress (Wilckens 1995; Sapolsky et al., 2000; Xu et al., 2009; Costantini et al., 2011).

While glucocorticoids have well-characterized immunosuppressive effects (Claman, 1972; De Bosscher et al., 2000), acute glucocorticoid release also enhances immune function through alterations in white blood cells and cytokine production (Dhabhar and McEwen, 1996; McInnis et al., 2015). As glucocorticoid elevations persist, however, the physiological response shifts to prevent long-term hyperactivity, resulting in decreased white blood cell counts and dysregulated expression of pro-inflammatory cytokines including interleukin-6 (IL-6) and tumor necrosis factor-α (TNF-α; Claman, 1972; Dhabhar, 2009). This highlights the role of GR-mediated glucocorticoid signaling in inflammation (Baschant and Tuckermann, 2010) and the opposing impacts of acute and chronic glucocorticoid exposure on immune function (Dhabhar and McEwen, 1997; Dhabhar, 2009). While both glucocorticoids and white blood cells play important roles in the production of reactive oxygen species (ROS; Smith and Weidemann, 1993; Yang et al., 2013; Costantini et al., 2011; Spiers et al., 2015), the contrasting actions of GR signaling with stressor duration obscure broad predictions of the role of HPA axis activation on ROS production and redox balance.

ROS are essential for cellular signaling and the immune response (Dröge, 2002; Hamanaka and Chandel, 2010; Yang et al., 2013), but dysregulated ROS generation promotes oxidative stress (Sies, 2019). White blood cells such as neutrophils (N) use superoxide and hydrogen peroxide generated during phagocytosis as part of the respiratory burst, an essential component of the innate immune response (Babior, 1984; Alberts et al., 2008). Moreover, mitochondrial and NADPH oxidase-derived ROS generation increase in response to GR signaling (McIntosh and Sapolsky, 1996; Houstis et al., 2006; You et al., 2009; Spiers et al., 2015). In addition to modulating ROS generation, glucocorticoids have differential impacts on antioxidants (Costantini, 2011). Acute glucocorticoid exposure increases antioxidant enzymes expression and activity (Yoshioka et al., 1994; Atanasova et al., 2009) while chronic exposure has an opposite effect (Djordjevic et al., 2010). The three-way interaction between glucocorticoids, immune cells, and ROS generation/removal thus complicates extrapolation of the impact of HPA axis activation on redox balance in animals with extreme life history events (Stier et al., 2019; Gormally and Romero, 2020; Ensminger 2021).

Northern elephant seals (*Mirounga angustirostris*) molt and develop during prolonged terrestrial fasts and frequently experience sleep apnea, hypoxemia, and ischemia/reperfusion events (Vázquez-Medina et al., 2012; Allen and Vázquez-Medina, 2019). In many animals, these processes increase oxidative stress and inflammation (Sakamoto et al., 1991; Colominas-Ciuró et al., 2019). Elephant seals, however, can sustain these fasts for months without experiencing oxidative stress or inflammation (Vázquez-Medina et al., 2010; Vázquez-Medina et al., 2013). Fasting-induced increases in antioxidants (Vázquez-Medina et al., 2010; Vázquez-Medina et al., 2011a) allow elephant seals to cope with physiological oxidative stress (Vázquez-Medina et al., 2012; Vázquez-Medina et al., 2013). Whether this adaptation of the redox system extends to the systemic response to HPA axis activation is unknown.

The impacts of glucocorticoids on redox homeostasis in marine mammals are poorly understood. Therefore, we studied whether acute or repeated activation of the HPA axis and GR signaling regulates immune cell function and redox balance in elephant seals. Elephant seals are a unique marine mammal species in which to study the impacts of acute and chronic HPA axis activation as they fast on land for months (Le Boeuf et al., 1973), maintain a functioning HPA axis response (Ensminger et al., 2014; McCormley et al., 2018), and are highly tractable which allows repeated sampling. Previous work in this species shows that HPA axis activation with exogenous ACTH increases circulating cortisol and aldosterone (Ensminger et al., 2014; McCormley et al., 2018). Additionally, *ex vivo* and *in vivo* transcriptomics studies (Khudyakov et al., 2015; Khudyakov et al., 2017; Deyarmin et al., 2019; Torres-Velarde et al., 2021) highlight the impacts of glucocorticoids on expression of genes involved in redox metabolism including Polo-like kinase 3, Thioredoxin, DNA damage inducible transcript 4, and Glutathione peroxidase (GPx) 4. Here, we used exogenous ACTH and a GR blocker to study the effects of acute and chronic HPA axis activation and GR signaling on oxidative stress and immune function in elephant seals. We found that ACTH increases adrenocorticosteroids and alters immune cell composition, and that GR signaling impacts aldosterone and white blood cell proportions. Unexpectedly, neither acute nor repeated activation of the HPA axis or GR signaling had an effect on oxidative stress or inflammatory cytokines, underscoring the robustness of elephant seals to sustained glucocorticoid elevations.

## 2. Methods

### 2.1 Study site and study animals

All animal procedures were approved by the Sonoma State University and UC Berkeley Institutional Animal Care and Use Committees and were conducted under the National Marine Fisheries Service permit # 19108. Sixteen post-weaned (simultaneously fasting and developing) early fasting (1-2 weeks, pre-molted) elephant seal pups were studied at Año Nuevo State Park, CA, USA.

### 2.2 Field procedures and sample collection

Animals were chemically immobilized with an intramuscular injection of ∼1 mg kg^-1^ tiletamine/zolazepam HCl (Telazol, Fort Dodge Animal Health, Fort Dodge, IA). Immobilization was maintained with intravenous injections of ketamine HCl (Ketaset, Fort Dodge Animal Health, Fort Dodge, IA). Pups were randomly assigned to one of three groups: 1) daily intramuscular injection of slow-release adrenocorticotropin LA gel (ACTH; Westwood Pharmacy, Richmond, VA) for four days, 2) subcutaneous implant of four-day time-release glucocorticoid receptor blocker pellets (RU486, Arcos Organics, Fair Lawn, NJ, Innovative Research of America, Sarasota, FL), or 3) both ACTH+RU486 treatment. Animals were given 0.22±0.01 U/kg (mean±SE) of ACTH and/or 3.130±0.102 mg/kg of RU486. ACTH doses were chosen based on previously published work from juvenile elephant seals (McCormley et al., 2018). RU486 implants were positioned laterally, approximately 220 mm superior to the pelvic girdle at the muscle interface, after making a small incision on the skin with a sterile scalpel and removing a blubber core using a 6.0 mm diameter biopsy punch (Vázquez-Medina et al., 2010). ACTH injections were given on the opposite side to where the biopsy was obtained. Body mass was collected via the truncated cones method (Crocker et al., 2001), which provides estimates within 5% of that obtained via direct measurement (Crocket et al., 2012). Blood samples were collected from the extradural vein into chilled serum and ethylenediaminetetraacetic acid (EDTA) vacutainer tubes for analysis at baseline and two hours post treatment. Four days later, animals were again immobilized and sampled two hours after the last ACTH injection (Fig 1). Samples were transported on ice to the laboratory where serum and plasma were prepared and stored at −80°C until analysis.

**Figure 1.**
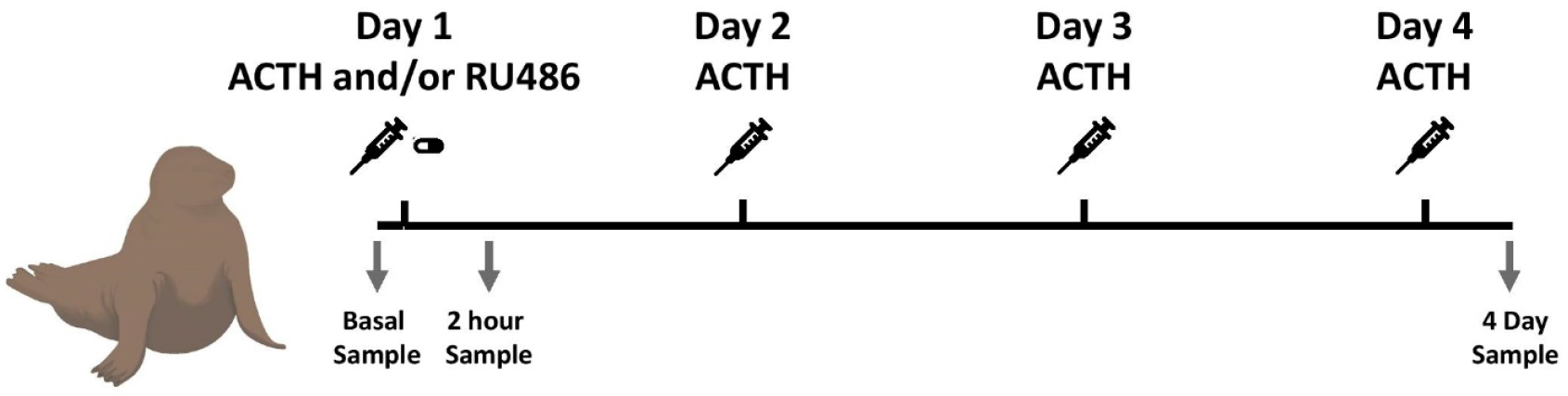
Field procedures and sample collection: Graphical representation of the experimental design for ACTH, and glucocorticoid receptor blocker (RU486) treatment and blood sampling. Seals were given either ACTH and/or RU486. Slow-release ACTH gel was administered every 24 hours for four days. A four-day time-release tablet of RU486 was implanted on day one. Blood samples were taken before treatment administration, two hours post-treatment administration, and two hours after the fourth day of treatment.

### 2.3 Whole blood, serum and plasma analysis

#### Hematology

Complete white blood cell counts were measured in whole blood using an automated hematology analyzer previously used to analyze blood collected from other pinnipeds (VetScan HM5, Abaxis Inc., Union City, CA; Unal et al., 2018, Thompson and Romano, 2019).

#### Corticosteroids and Metabolites

Cortisol (11-CORHU-E01, Alpco, Salem, NH) and aldosterone (11-AD2HU-E01, Alpco, Salem, NH) were measured in serum using EIA kits validated for use in elephant seals (McCormley et al., 2018). Non-esterified fatty acids (NEFA; HR Series NEFA-HR(2), Wako Chemicals, Richmond, VA), and triglycerides (10010303, Cayman Chemical, Ann Arbor, MI) were measured in plasma using enzymatic colorimetric assays as previously described (Ortiz et al., 2003; Viscarra et al., 2012).

#### Antioxidants

Plasma GPx (703102, Cayman Chemical, Ann Arbor, MI), glutathione-disulfide reductase (GSR; 703202, Cayman Chemical, Ann Arbor, MI), and catalase activities (707002, Cayman Chemical, Ann Arbor, MI), along with total thiols (700340, Cayman Chemical, Ann Arbor, MI) were measured using colorimetric kits previously used in elephant seals (Vázquez-Medina et al., 2010; Sharick et al., 2015), or validated via parallelism and spiked recovery (total thiols).

#### Oxidative damage

Plasma F_2_-isoprostanes, a marker for lipid peroxidation, were measured using gas chromatography-mass spectrometry at the Vanderbilt University Eicosanoid Core Laboratory as previously described (Milne et al., 2007; Vázquez-Medina et al., 2010). Protein carbonyls were measured using commercial EIA assays (STA-310, Cell Bio Labs, San Diego, CA) validated for elephant seals via parallelism and spiked recovery.

#### Cytokines

Plasma transforming growth factor beta (TGF-β; DY240, R&D Systems, Minneapolis, MN), and IL-6 (ELC-IL6-1, RayBioTech, Norcross, GA) were measured using commercial kits previously validated for elephant seals (IL-6; Peck et al. 2015) or validated via parallelism and spiked recovery (TGF-β).

All samples were analyzed in duplicate. The average coefficient of variation for blood, plasma, and serum analyses was 6.33% for intra-assay and 3.84% for inter-assay variation.

### 2.4 Data analysis

Data were analyzed using linear mixed models (v.1.1.463, R Development Core Team, Boston, MA; package: lme4) and met the assumptions of the models. Figures were made in RStudio (package: ggplot2). For all models, treatment, time point, and the interaction of treatment and time point were included as fixed effects and seal ID was included as a random effect.

Interaction terms were removed if they did not significantly explain variation in the models (*p* > 0.20). Seal mass and mass-specific treatments were originally included in the models but were removed as they did not explain variation. Tukey’s post hoc tests were used to identify specific effects. Line graph data are represented as mean ± standard error of mean and box and whisker data are presented as median, upper quartile, lower quartile, and 1.5*interquartile range (whiskers). Results were considered significant at *p* < 0.05.

## 3. Results

### 3.1 Repeated activation of the HPA axis causes differential effects on adrenal steroids

We measured cortisol and aldosterone to corroborate acute and sustained activation of the HPA axis by exogenous ACTH and RU486. Both treatment and time had an effect on serum cortisol and aldosterone (cortisol: F_2,52_=42.549, p<0.001; F_2,52_=220.069, p<0.001; aldosterone: F_2,52_=4.880, p=0.023; F_2,52_=57.087, p<0.001). There was an interaction between treatment and time which led to differential effects on circulating cortisol and aldosterone (F_4,50_=52.187, p<0.001; F_4,50_=8.775, p<0.001). Acute ACTH injection increased both cortisol and aldosterone after two hours (Fig 2A, 1B). Similarly, both the cortisol and aldosterone post-ACTH responses were maintained after four daily ACTH injections; however, the magnitude of the cortisol response at four days was attenuated compared to two hours (Fig 2A, 2B). ACTH+RU486 did not change the response compared to ACTH alone. Moreover, aldosterone increased in response to RU486 alone at four days (Fig 2A, 2B). These results show that 1) exogenous ACTH administration increases adrenocorticosteroids in post-weaning elephant seals, 2) repeated ACTH treatment for four days decreases the cortisol but not the aldosterone response, and 3) GR blockade with RU486 for four days causes a compensatory increase in circulating aldosterone.

**Figure 2.**
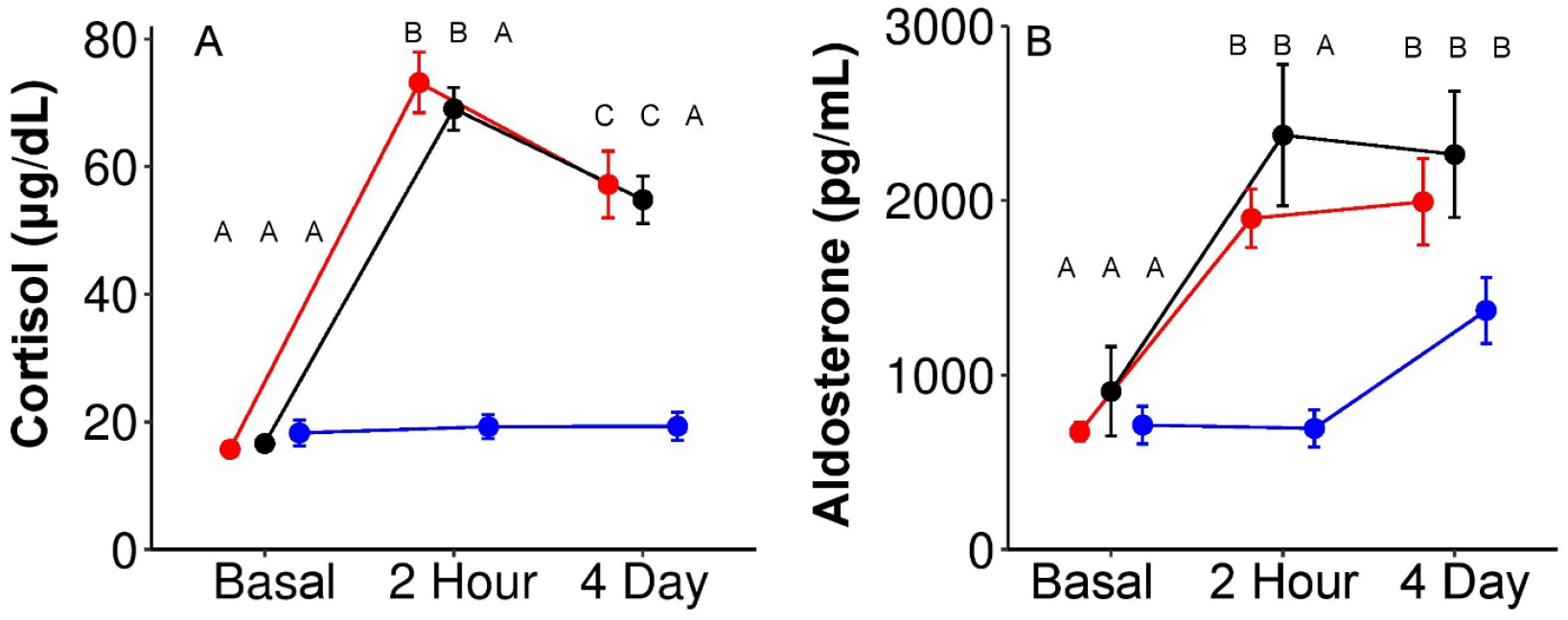
Repeated HPA axis activation differentially affects cortisol and aldosterone: Serum adrenocorticosteroid levels during basal (pre-treatment) conditions and after two hours and four days of treatment. A) Cortisol and B) aldosterone. ACTH: red (n=6), ACTH+RU486: black (n=6), RU486 alone: blue (n=6). Different letters represent statistical differences of the interaction of time and treatment respectively (p<0.05) based on Tukey’s post-hoc comparisons of linear mixed models. Data are mean ± standard error of mean.

### 3.2 Repeated stimulation of the HPA axis does not induce oxidative stress in fasting elephant seal pups

Sustained glucocorticoid release induces oxidative stress in a variety of vertebrates (Costantini et al., 2011). Hence, we measured circulating antioxidants, lipid peroxidation and protein oxidation to assess the impact of acute and repeated HPA stimulation on oxidative stress in fasting elephant seal pups. While treatment did not alter F_2_-isoprostanes, there was an impact of time on F_2_-isoprostanes (F_2,52_=2.697, p=0.100; F_2,52_=4.612, p=0.017), with concentrations lower at four days than at two hours (Fig 3A). There was no impact of treatment or time on protein carbonyls (F_2,52_=0.266, p=0.770; F_2,52_=2.599, p=0.089; Fig 3B). Together, these data show that neither acute nor repeated stimulation of the HPA axis or endogenous GR blockade for four days causes systemic oxidative damage in fasting elephant seal pups.

**Figure 3.**
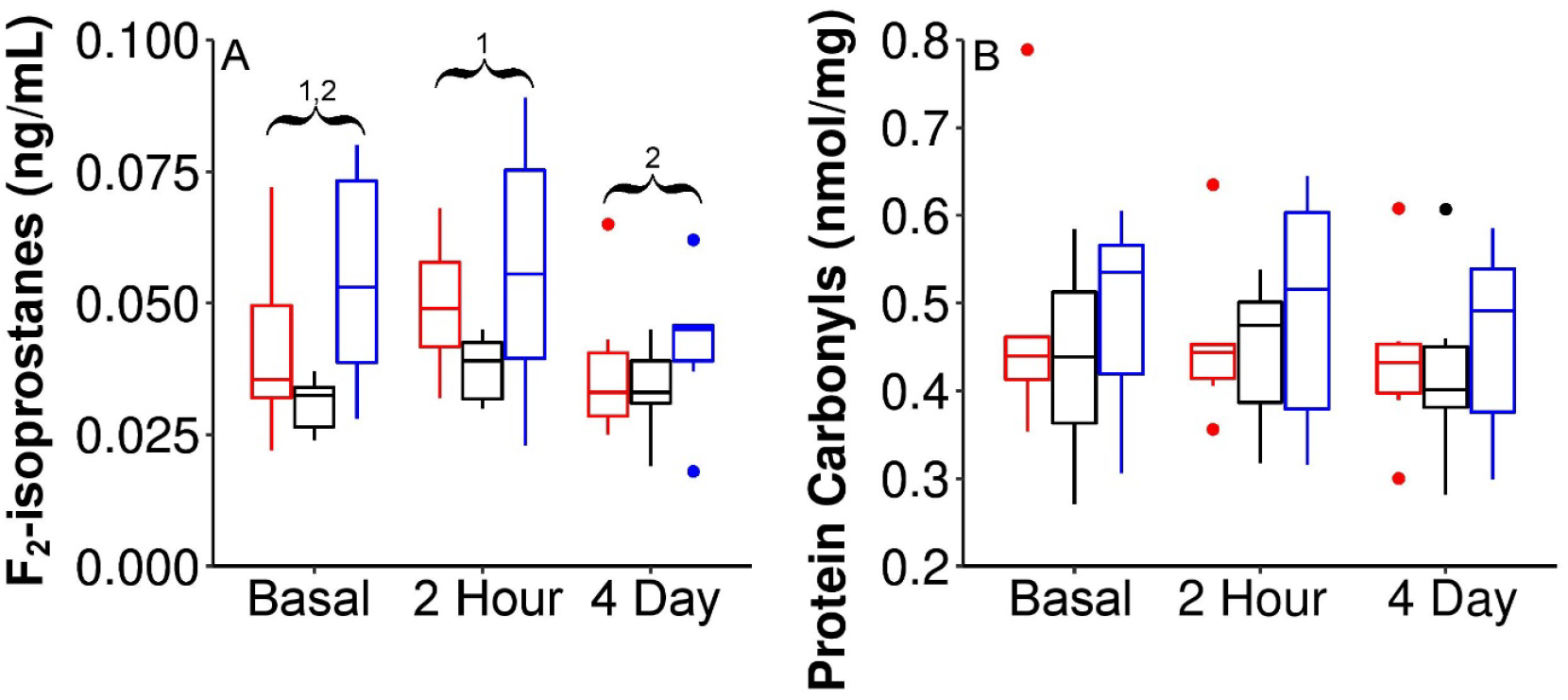
Neither acute nor repeated HPA axis activation nor GR blockade alter oxidative damage: Plasma oxidative damage markers during basal conditions and after two hours and four days of treatment. A) F_2_-isoprostanes and B) protein carbonyls. ACTH: red (n=6), ACTH+RU486: black (n=6), RU486 alone: blue (n=6). Different numbers represent statistical differences between time groups (p<0.05) based on Tukey’s post-hoc comparisons of linear mixed models. Boxplots depict the first quartile and third quartile plus (box) ± 1.5*interquartile range (whiskers) and the median (horizontal line).

We then measured plasma antioxidants to explore whether manipulation of the HPA axis or GR blockade alters antioxidant defenses in fasting elephant seal pups. Treatment did not impact catalase or GSR activity (F_2,52_=0.669, p=0.528; F_2,52_=0.585, p=0.570). Surprisingly, both catalase and GSR activity decreased at four days compared to baseline (F_2,52_=9.705, p<0.001; F_2,52_=5.113, p=0.011; Fig 4 A, B). Treatment did not impact GPx, but, in contrast to catalase and GSR, GPx activity increased at four days compared to baseline and two hours (F_2,52_=0.602, p=0.561; F_2,52_=6.921, p=0.003; Fig 4C). There were trends for the impact of treatment and time on total thiols (F_2,52_=3.225, p=0.069; F_2,52_=2.913, p=0.070). RU486 tended to decrease total thiols compared to ACTH+RU486, while concentrations tended to be higher on day four compared to two hours (Fig 4D). There was no interaction between treatment and time for total thiols (F_4,50_=1.946, p=0.129). These data show that increased antioxidant defenses do not account for the absence of systemic oxidative damage during acute or repeated stimulation of the HPA axis or endogenous GR blockade in elephant seals. Moreover, these data show that fasting progression is associated with increased GPx activity and decreased lipid peroxidation.

**Figure 4.**
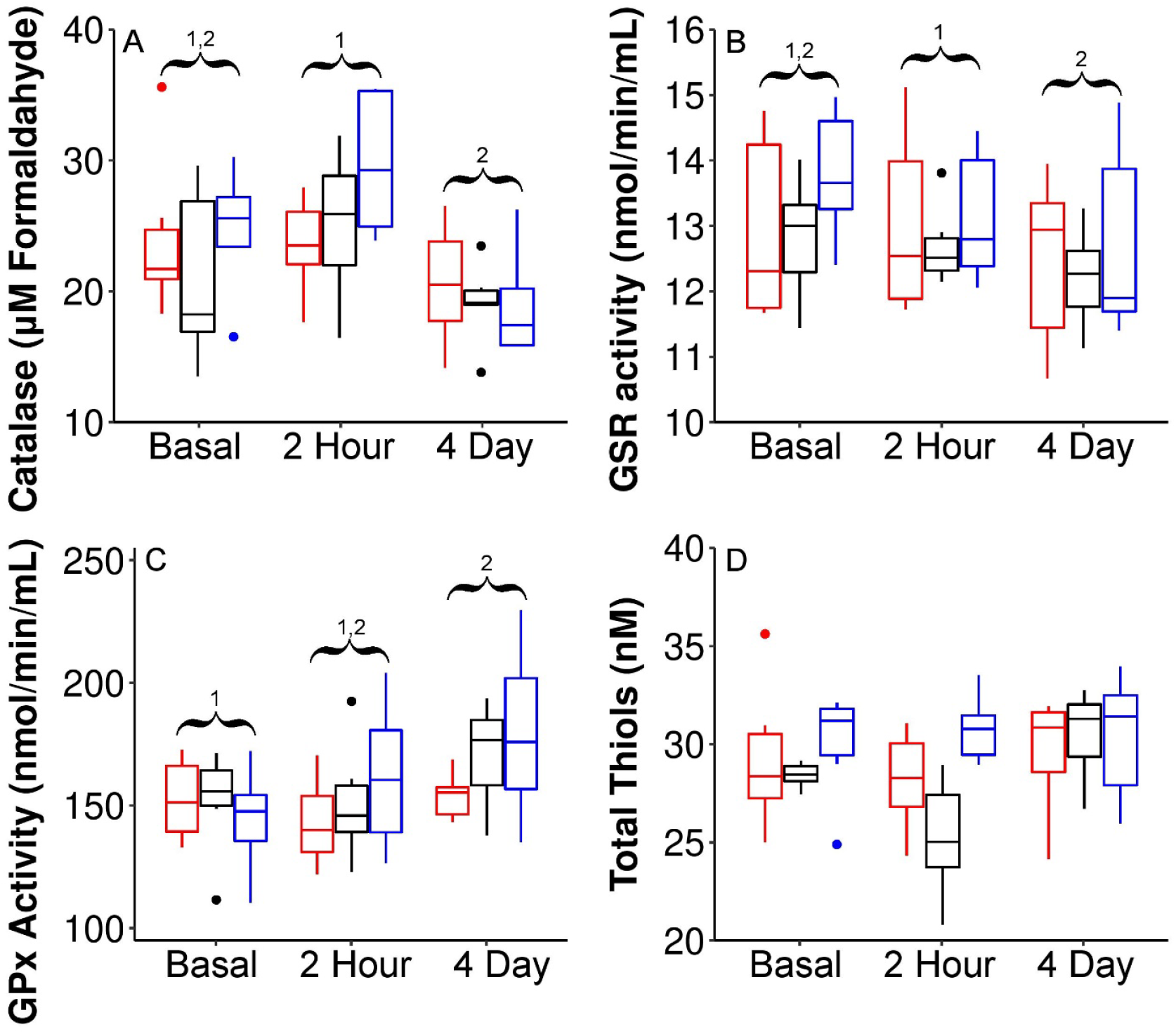
Neither acute nor repeated HPA axis activation nor GR blockade alter circulating antioxidants: Plasma antioxidants during basal conditions and after two hours and four days of treatment. A) Catalase activity, B) glutathione-disulfide reductase (GSR) activity, C) glutathione peroxidase (GPx) activity, and D) total thiols. ACTH: red (n=6), ACTH+RU486: black (n=6), RU486 alone: blue (n=6). Different numbers represent statistical differences between time groups (p<0.05) based on Tukey’s post-hoc comparisons of linear mixed models. Boxplots depict the first quartile and third quartile plus (box) ± 1.5*interquartile range (whiskers) and the median (horizontal line).

### 3.3 Stimulation of the HPA axis does not induce lipolysis in postweaning elephant seals

In most animals, glucocorticoid release increases fuel availability through the liberation of stored energy. Hence, we measured circulating NEFA and triglyceride levels to assess the impact of acute and repeated stimulation of the HPA axis on circulating lipids. NEFA was not altered by treatment, time, or the interaction of treatment and time (F_2,52_=0.663, p=0.530; F_2,52_=0.771, p=0.471; F_4,50_=2.144, p=0.100; Table 1). Similarly, triglycerides were not affected by treatment or the interaction of treatment and time (F_2,52_=0.630, p=0.546; F_4,50_=2.135, p=0.101). However, triglycerides were impacted by time, with triglyceride concentrations being lower at four days compared to baseline or two hours (Table 1). These data show that neither ACTH nor GR signaling alter rates of lipolysis in post-weaned (simultaneously fasting and developing) elephant seals.

**Table 1.** Plasma lipids and cytokines in elephant seals pups during basal conditions and after ACTH and RU486 administration. * denotes significant differences from baseline (p<0.05) based on Tukey’s post-hoc comparisons of linear mixed models.

|  | ACTH |  |  | ACTH+RU486 |  |  | RU486 |  |  |
| --- | --- | --- | --- | --- | --- | --- | --- | --- | --- |
|  | Basal | 2 hour | 4 day | Basal | 2 hour | 4 day | Basal | 2 hour | 4 day |
| NEFA (mmol/L) | 1.56±0.21 | 1.55±0.25 | 1.66±0.22 | 1.24±0.09 | 1.94±0.44 | 1.2±0.14 | 1.19±0.13 | 1.14±0.12 | 1.61±0.38 |
| Triglycerides (mg/mL) | 21.46±4.45 | 21.37±4.13 | 11.6±1.44* | 15.45±0.96 | 18.19±2.81 | 10.39±1.3* | 16.75±1.92 | 15.45±1.29 | 17.46±4.7* |
| TGF-β (pg/mL) | 146±31.5 | 163.6±16.2 | 121.5±22.3 | 85.2±9.3 | 150.5±32.6 | 121.1±23.5 | 185.5±35.8 | 227.6±54.1 | 136.3±56.6 |
| IL-6 (ng/mL) | 0.113±0.041 | 0.141±0.02 | 0.116±0.027 | 0.041±0.011 | 0.135±0.029 | 0.088±0.032 | 0.149±0.02 | 0.147±0.027 | 0.072±0.033 |

### 3.4 Acute and repeated stimulation of the HPA axis alters white blood cell counts without affecting cytokine levels

We measured white blood cell counts and cytokine levels to examine the role of acute and repeated stimulation of the HPA axis and GR signaling on the immune system in elephant seals. Neither treatment nor time altered total white blood cell counts (F_2,52_=0.238, p=0.791; F_2,52_=0.432, p=0.653). There was a trend for an interaction, however, post-hoc analysis revealed no differences (F_4,50_=2.607, p=0.057; Fig 5A). Treatment did not affect N%, but N% increased at four days compared to baseline and two hours (F_2,52_=2.625, p=0.108; F_2,52_=7.984, p=0.001 Fig 5B). In contrast, both treatment and time impacted lymphocyte (L) % (F_2,52_=4.433, p=0.032; F_2,52_=11.070, p<0.001). ACTH lowered L% compared to ACTH+RU486 or RU486, and L% was also lower at four days than either baseline or at two hours (Fig 5C). Similar to L%, both treatment and time altered the N:L ratio (F_2,52_=5.927, p=0.014; F_2,52_=13.132, p<0.001). ACTH increased the N:L ratio compared to ACTH+RU486 or RU486. The N:L ratio was higher at four days compared to baseline and two hours (Fig 5D). These results show that acute and repeated stimulation of the HPA axis have differential effects on white blood cell counts in elephant seals. While acute stimulation of the HPA axis does not have an effect on the N:L ratio, repeated activation increases N:L suggesting a shift in immune response mediated by GR.

**Figure 5.**
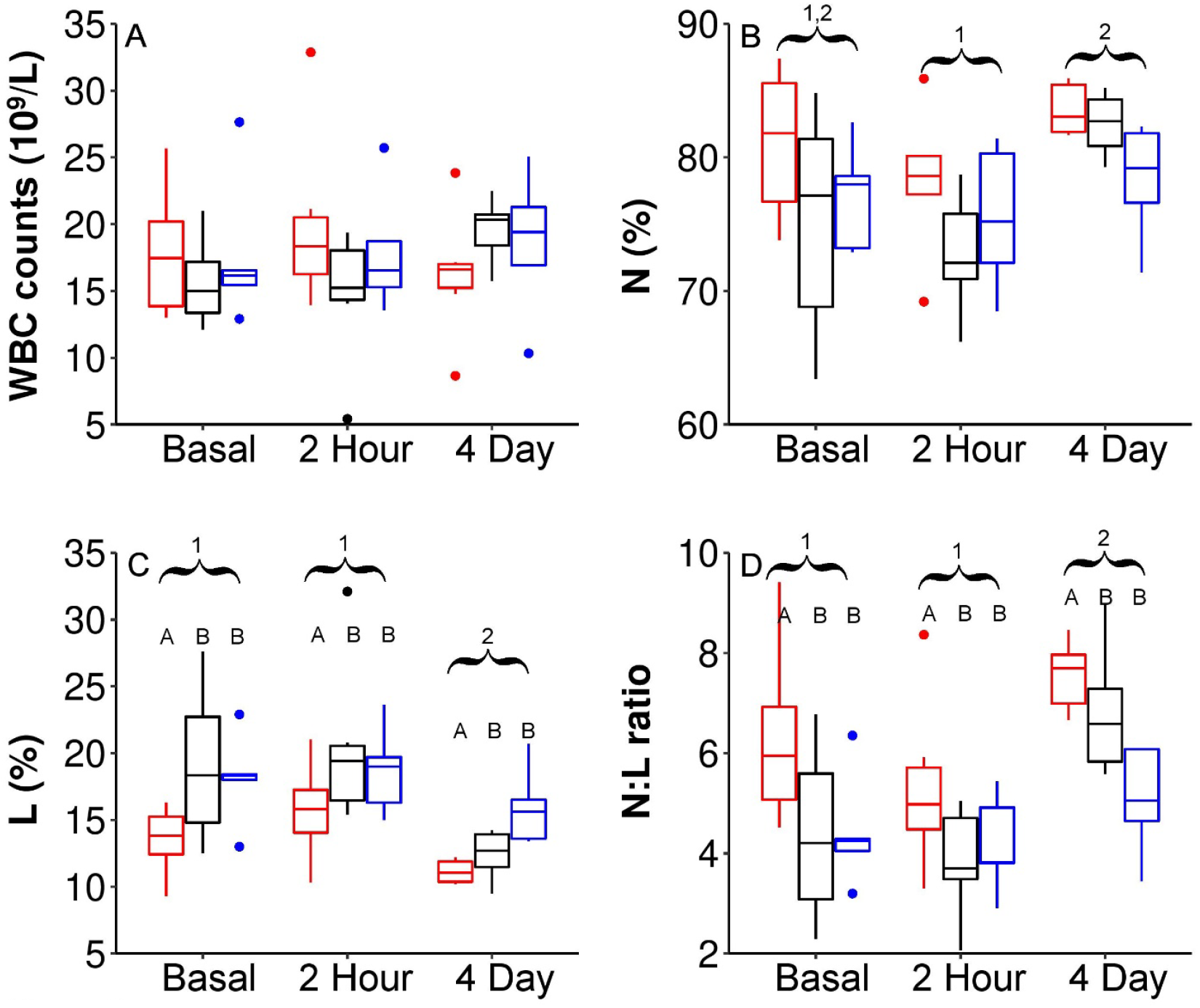
Acute and repeated activation of the HPA axis alters white blood cell counts: White blood cell counts and composition during basal conditions and after two hours and four days of treatment. A) Complete white blood cell (WBC) counts, B) neutrophil percentage (N), C) lymphocyte percentage (L), and D) neutrophil to lymphocyte (N:L) ratio. ACTH: red (n=6), ACTH+RU486: black (n=6), RU486 alone: blue (n=6). Different numbers represent statistical differences between time groups (p<0.05) and different letters represent statistical differences between treatment groups based on Tukey’s post-hoc comparisons of linear mixed models. Boxplots depict the first quartile and third quartile plus (box) ± 1.5*interquartile range (whiskers) and the median (horizontal line).

We then measured plasma cytokine levels to explore whether HPA axis activation or GR signaling alters circulating cytokine levels. Neither treatment nor time altered TGF-β or IL-6 levels (TGF-β: F_2,52_=2.112, p=0.155; F_2,52_=2.640, p=0.086; IL-6: F_2,52_=1.404, p=0.276; F_2,52_=2.632, p=0.087; Table 1). These data show that despite altering white blood proportions, repeated stimulation of the HPA axis does not increase cytokine expression in elephant seals, underscoring these animals’ extraordinary resistance to inflammation while simultaneously fasting and developing.

## 4. Discussion

A variety of stressors activate the HPA axis, altering corticosteroid concentrations, redox balance, and inflammation across taxa. While the stress response has been heavily explored in terrestrial vertebrates, considerably less is known about the downstream impacts of this response in marine mammals. Here we show that acute and chronic activation of the HPA axis elicit differential responses on adrenocorticosteroids and that GR blockade causes a compensatory increase in aldosterone but not cortisol in elephant seal pups. Additionally, we found that neither manipulation of the HPA axis nor blockade of endogenous GR signaling induces oxidative stress or lipolysis in post-weaned elephant seal pups, though both are impacted as the fast progresses. Our data also show that stimulation with exogenous ACTH alters the proportion of WBC without altering cytokine levels, and that this effect is driven by GR signaling.

In our experiments, ACTH stimulation increased circulating cortisol and aldosterone, however, the impacts of chronic ACTH exposure and the role of GR feedback differed between the two adrenocorticosteroids. The magnitude of the cortisol response to ACTH stimulation decreased after four days of repeated ACTH exposure, as reported previously in juvenile elephant seals (McCormley et al., 2018). This response may result from desensitization of the HPA axis, stronger negative feedback, or adrenal exhaustion (Rich and Romero, 2005; Welberg et al., 2006; Gądek-Michalska et al., 2013). HPA axis desensitization was previously observed in adult male elephant seals (Ensminger et al., 2014), suggesting that repeated activation of the HPA axis results in desensitization of the HPA axis independent of age.

Our results also show low individual variability in cortisol levels in response to ACTH in elephant seal pups, suggesting tight regulation of cortisol during early postnatal development. In contrast, the cortisol response to ACTH is largely variable in juvenile and adult elephant seals (Ensminger et al., 2014; McCormley et al., 2018), suggesting that this variability is impacted by postnatal maturation. Our results about the effect of ACTH stimulation on aldosterone are consistent with those of juvenile elephant seals (McCormley et al., 2018), but contrary to adult males, suggesting that life history stage impacts negative feedback on aldosterone secretion (Ensminger et al., 2014). Contrary to cortisol, there was a compensatory impact of GR blockade on aldosterone after four days of repeated ACTH stimulation, further supporting the hypothesis that aldosterone is an important component of the stress response in marine mammals (Thomson and Geraci, 1986; Houser et al., 2011). Though chronically elevated aldosterone may have less impact on land due to low urinary output (Adams and Costa, 1993), it could negatively impact osmotic balance while at sea due to aldosterone’s influence on sodium and potassium concentrations (Morris, 1981; Ortiz et al., 2000). These differential impacts on aldosterone and cortisol also show the differential sensitivity of the zona glomerulosa and zona fasciculata to acute and repeated ACTH stimulation and endogenous GR blockade.

Despite robust HPA axis activation, which elevated both cortisol and aldosterone, we found no effects of ACTH stimulation or GR blockade on oxidative stress in fasting elephant seal pups. Cortisol increases oxidative stress in several vertebrates (Costantini et al., 2011; Spiers et al., 2015) and aldosterone induces oxidative stress via activation of NADPH oxidases (Sun et al., 2002; Miyata et al., 2005); however, our results show that neither cortisol/aldosterone nor GR signaling regulate circulating antioxidants, lipid peroxidation or protein oxidation in elephant seal pups. These results support previous observations of the extraordinary capacity elephant seals possess to cope with oxidative stress derived from prolonged food and water deprivation (Vázquez-Medina et al., 2010; Vázquez-Medina et al., 2011a; Vázquez-Medina et al., 2013), sleep apnea, hypoxemia, and ischemia/reperfusion (Vázquez-Medina et al., 2012; Allen and Vázquez-Medina, 2019). Both acute and repeated ACTH stimulation alter blubber and muscle expression of genes involved in redox balance in juveniles (Polo-like kinase 3 and Thioredoxin; Khudyakov et al., 2015; Khudyakov et al., 2017; Deyarmin et al., 2019). Similarly, sustained exposure to glucocorticoids upregulates the expression of the phospholipid hydroperoxidase GPx 4 while downregulating peroxiredoxin 6 expression in elephant seal muscle cells in primary culture (Torres-Velarde et al., 2021). Therefore, the lack of response here suggests that either pups have an altered oxidative response to cortisol compared to other life history stages, likely due to the combination of fasting and development, or that the effect of cortisol on oxidative stress is tissue-specific. In support of the latter hypothesis, a previous meta-analysis shows tissue-specific differences in cortisol-induced oxidative stress across taxa (Costantini et al., 2011). Future research should focus on identifying the interplay between tissue-specific and systemic oxidative responses to further elucidate these relationships.

While our treatments did not alter circulating antioxidant enzymes or oxidative damage, we found opposing patterns in lipid peroxidation and antioxidants involved in reducing lipid hydroperoxides (GPX and total thiols) within the short (four day) fasting progression. Lipid peroxidation decreased from day one to day four and was not associated with changes in NEFA or triglyceride levels (data not shown) (Pérez-Rodríguez et al., 2015). Moreover, GPx and total thiols increased with time, supporting the hypothesis that fasting promotes a positive redox balance in part by stimulating the glutathione system, as previously shown in fasting elephant seal pups (Vázquez-Medina et al., 2010; Vázquez-Medina et al., 2011a; Ensminger et al., 2021). Interestingly, both catalase and GSR decrease from two hours to four days. Combined with the increase in GPx and total thiols, these data suggest a potential shift in resources away from recycling glutathione through GSR and toward increasing lipid hydroperoxide removal and stimulating glutathione synthesis (Vázquez-Medina et al., 2011a). Of note, the observed decrease in catalase and GSR opposed patterns previously found in muscle and red blood cells, further suggest tissue-specific effects in redox metabolism or differential effects of short and long fasting duration (four days vs two months; Vázquez-Medina et al., 2010; Vázquez-Medina et al., 2011a).

Similar to oxidative stress markers, we found no impact of ACTH or GR blockade on NEFA or triglycerides; however, triglycerides decreased with fasting progression. While changes in triglycerides may represent a fasting-derived decrease in stored fat supplies (Williams et al., 1999), the constant NEFA concentrations suggest that these seals were not fat limited (Jenni-Eiermann and Jenni, 1992). As elephant seals have a high fat-based metabolism, tight regulation of cortisol-induced lipolysis may support fat metabolism during prolonged fasting (Crocker et al., 2014). However, while breeding females show correlations between cortisol and NEFA, males only exhibit this relationship during the molt (Ensminger et al., 2014, Fowler et al., 2016). As such, our results suggest that life history plays a strong role in the downstream effects of HPA axis activation and cortisol on lipolysis in elephant seals and that animals in certain life history stages (weaned elephant seal pups and breeding males) might limit the impact of adrenal stimulation on fat metabolism during prolonged fasts.

While there was no impact on oxidative balance or lipolysis, repeated ACTH stimulation shifted the composition of white blood cells, lowering the L proportion and subsequently increasing the N:L ratio. While white blood cells take longer to respond to stressors (Gross, 1990; DuRant et al., 2015), they stay elevated over the duration of chronic stress exposure (Goessling et al., 2015). When combined with findings that changes in white blood cells are more sensitive than cortisol to a wider range of stressors (Müller et al., 2011), these results support the hypothesis that N:L ratios may be a better marker for assessing chronic stress exposure in wildlife, and that the observed changes in N:L ratio are likely driven by both N and L (Keogh and Atkinson, 2015; Davis and Maney, 2018). Despite shifts in white blood cell proportions and cortisol, we found no impact of either ACTH stimulation or GR blockade on pro-inflammatory cytokines. Similar cytokines remain stable across the fast despite increases in cortisol (i.e., IL-1β; Ortiz et al., 2003; Vázquez-Medina et al., 2013; Peck et al., 2016), suggesting that elephant seals possess physiological mechanisms to limit inflammation. This is further supported by the intrinsic anti-inflammatory properties of elephant seal serum (Bagchi et al., 2018), though more research is needed to understand the uncoupling of both cortisol and HPA axis activation from the inflammatory response, as activation of the renin-angiotensin-aldosterone system increases with fasting progression in elephant seal pups, along with muscle TNF-α muscle expression and protein abundance (Suzuki et al., 2013; Vázquez-Medina et al., 2010; Vázquez-Medina et al., 2013).

Overall, this study shows that neither repeated stimulation of the HPA axis nor blockade of endogenous GR signaling causes systemic oxidative stress, inflammation or alters lipolysis in simultaneously fasting and developing weaned elephant seals. Hence, these results underscore elephant seals’ robust tolerance of repeated and sustained cortisol elevations. Furthermore, our results support the hypothesis that animals in metabolically demanding life history stages or living in areas with repeated stressors may rely on physiological processes that help mitigate the deleterious impacts of a prolonged stress response (Huber et al., 2017; Stier et al., 2019; Ensminger et al., 2021). These data also support the hypothesis that aldosterone is an important component of the stress response in marine mammals and highlights a potential role of GR signaling in osmotic regulation. Though the sample size precludes an explicit examination of the interaction on treatment and fasting N:L ratio, these data highlight the need for future research to examine the use of this metric as a more consistent indicator of chronic stress in marine mammals (Davis and Maney et al., 2018).

## 5. Acknowledgements

We thank B. Gabriela Arango, Andrea Salvador-Pascual, Julia María Torres-Velarde, Anna Castello, Diamond Luong, Alex Li, and Marvin Miller for assistance with sample collection and the rangers at Año Nuevo State Park for assistance with logistics. DCE was supported by a National Science Foundation Postdoctoral Research Fellowship (grant #1907155). EKL was supported by a Department of Defense National Defense Science and Engineering Graduate Fellowship. KNA was supported by a National Science Foundation Graduate Research Fellowship (grant #1752814). Research funded by UC Berkeley startup funds. No competing interests declared.

